# “Multi-Agent” Screening Improves the Efficiency of Directed Enzyme Evolution

**DOI:** 10.1101/2021.04.06.438652

**Authors:** Tian Yang, Zhixia Ye, Michael D. Lynch

## Abstract

Enzyme evolution has enabled numerous advances in biotechnology. However, directed evolution programs can still require many iterative rounds of screening to identify optimal mutant sequences. This is due to the sparsity of the fitness landscape, which in turn, is due to “hidden” mutations that only offer improvements synergistically in combination with other mutations. These “hidden” mutations are only identified by evaluating mutant combinations, necessitating large combinatorial libraries or iterative rounds of screening. Here, we report a multi-agent directed evolution approach that incorporates diverse substrate analogues in the screening process. With multiple substrates acting like multiple agents navigating the fitness landscape, we are able to identify “hidden” mutant residues that impact substrate specificity without a need for testing numerous combinations. We initially validate this approach in engineering a malonyl-CoA synthetase for improved activity with a wide variety of non-natural substrates. We found that “hidden” mutations are often distant from the active site, making them hard to predict using popular structure-based methods. Interestingly, many of the “hidden” mutations identified in this case are expected to destabilize interactions between elements of tertiary structure, potentially affecting protein flexibility. This approach may be widely applicable to accelerate enzyme engineering. Lastly, multi-agent system inspired approaches may be more broadly useful in tackling other complex combinatorial search problems in biology.

**Highlights:**

- “Multi-agent” screening improves directed evolution.
- The incorporation of multiple substrates leads to the identification of “hidden” mutations, which can be hard to identify through one substrate.
- “Hidden” mutations are often remote from the active site and are expected to interrupt stabilizing side-chain interactions, thus increasing enzyme flexibility.

## Introduction

Enzyme engineering has led to exciting developments including novel enzymatic reactions thanks to the advances in directed evolution. ^1–3^ Additionally, enzyme engineering programs aimed at expanding the substrate utilization of known enzymes are more commonplace than ever. ^4,5^ These directed evolution strategies rely on first generating genetic diversity followed by screening and/or selection to identify genetic variants with improved performance, ^6,7^ and have become one of the most powerful and widespread tools in biological research with a wide range of applications.^8–11^

However, the directed evolution of enzymes and all proteins is still challenged by the sheer combinatorial complexity of the sequence design space (20^n^, n is the number of amino acids) as well as the sparsity of beneficial mutations in this fitness landscape.^12^ Over the past three decades, researchers have devised various methodologies to address these issues. ^13^ Structure-guided or targeted mutagenesis methods can prioritize residues and significantly reduce the size of the sequence space for laboratory evaluation, but this requires accurate structural information, and oftentimes subjective prioritization will miss unpredictable mutants and mutant combinations^14^. More objective structure guided approaches are often computational in nature.^15^ Newer machine learning assisted methods have emerged with an attractive potential to virtualize the screening process. ^16–18^ However, these methods often require intensive computational resources and/or training on large existing datasets.^17^ High-throughput screening methods, which enable the rapid evaluation of large libraries, are becoming more commonplace and new tools allowing for increases in throughput are always being developed. ^19–23^ However, even the fastest screening capabilities are still only able to evaluate a small fraction of the massive combinatorial search space(Figure S1) and as a result brute force screening can still be likened to finding a needle in a haystack.^24–27^

As mentioned, in the case of proteins and enzymes, the fitness landscape is extremely sparse.^12^ That is to say that a vast majority of mutants in the 20^n^ sequence space offer no benefit to a given target activity. A primary reason for the sparsity of these landscapes is the presence of “hidden” mutations. As defined here, these “hidden” mutations only offer a benefit synergistically (when in combination) with other mutation(s) and as a result are “hidden” in initial rounds of scanning mutagenesis (screening small libraries of sequences with single amino acid mutations Figure 1a). These mutations, by definition, can only be identified by either screening of large combinatorial libraries that could exceed the capacity of high-throughput screening, or iterative rounds of screening of smaller libraries^28^ that can lead to “fitness valleys”.^29^

**Figure 1.**
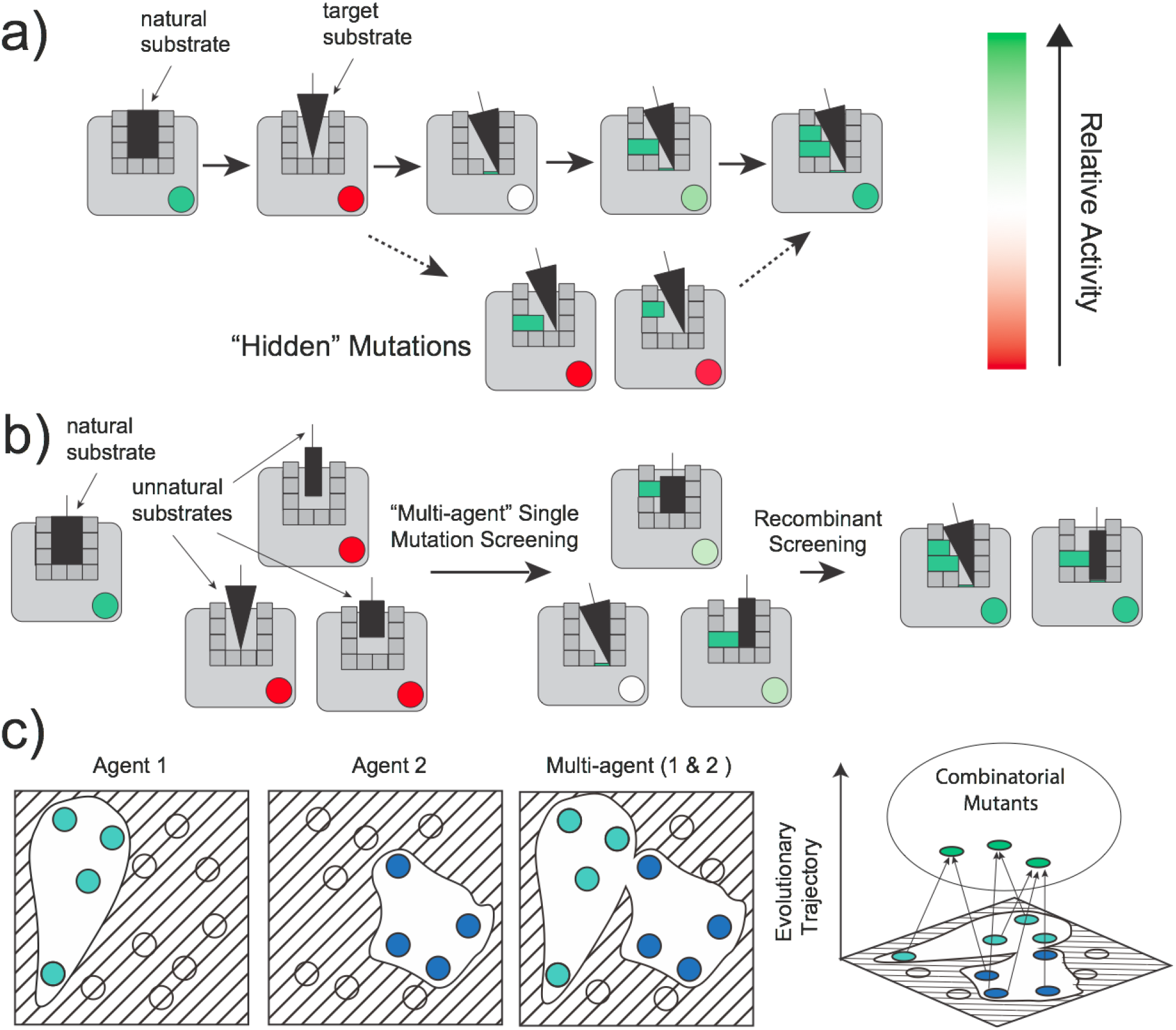
An overview of “hidden” mutations and multi-agent screening. **a:** Dark gray squares represent individual amino acids in the larger protein structure (light gray), substrates are illustrated in black. Beneficial amino acid mutations are shown in green, with the relative activity of the mutant overall, given by the colored circle. In the directed evolution of enzymes, “hidden” mutations are defined as mutations that only offer benefit in performance when in combination with other mutations and may be neutral or detrimental to activity alone. Such synergistic mutations can be hard to predict in rational and structural-guided methods, and can only be identified in the later iterations of mutation scanning; **b:** Incorporating multiple substrate analogues (black shapes) to identify “hidden” mutations. Mutations hidden for one substrate may have a positive effect for another. **c:** A multi-agent approach to navigate the mutant fitness landscape. Circles are mutants present in the optimal solution. Solutions visible to agent 1(light green) are not necessarily visible to agent 2 (blue) and vice versa. Incorporating results from both agents will lead to the more complete evaluation of solutions. Recombining solutions visible to each individual agent (green and blue) can lead to improved performance with greater efficiency.

In this work, we hypothesized that using multiple substrate analogues as multiple agents navigating the sequence space would increase the efficiency of searching for beneficial mutations. In computer science, multi-agent systems refer to systems consisting of multiple intelligent agents operating in a shared environment in a goal-oriented way. ^30,31^ Such systems can often tackle problems that are difficult or impossible to solve with single agents.^32^ Current directed enzyme evolution approaches can be viewed as single agent systems, where one agent (substrate) searches the vast sequence space for beneficial sequences for one predetermined property (activity). Hidden mutations can be thought of as “hidden” to this single agent in initial searching (Figure 1c). Multiple substrate analogues are like multiple agents (Figure 1b, 1c), where a “hidden” mutation impacting substrate specificity may be hidden for a single agent but not for another. These alternative agents may identify hidden mutant residues earlier in directed evolution trajectories with smaller sized libraries.

To test our hypothesis and demonstrate the utility of this approach, we chose to engineer the substrate specificity of malonyl-CoA synthetase (MatB) of *Rhizobium trifolii,* an enzyme known for its potential in the synthesis of natural products and natural product analogues. ^33^ As illustrated in Figure 2a, we simultaneously evolved MatB’s activity on 17 structural analogues of malonate, its native substrate. This screening identified numerous “hidden” mutations, with a total of 20 mutations found affecting activity. On the contrary, at most only 4 mutations were identified when considering only a single substrate. By screening a small combinatorial library constructed from this larger group of mutations we were able to identify a group of enzyme variants with significantly improved activity for 10 substrates. Notably, these variants have a defined specificity, and broadly active, nonspecific mutants were not identified. Lastly, we investigated the potential mechanism of action of the mutations we identified, particularly those “hidden mutations” distant from the active site. The majority of non active site mutations are predicted to destabilize interactions between different strands of the folded protein, which may enable increased protein flexibility and in combination with active site mutations altered substrate specificity.

**Figure 2.**
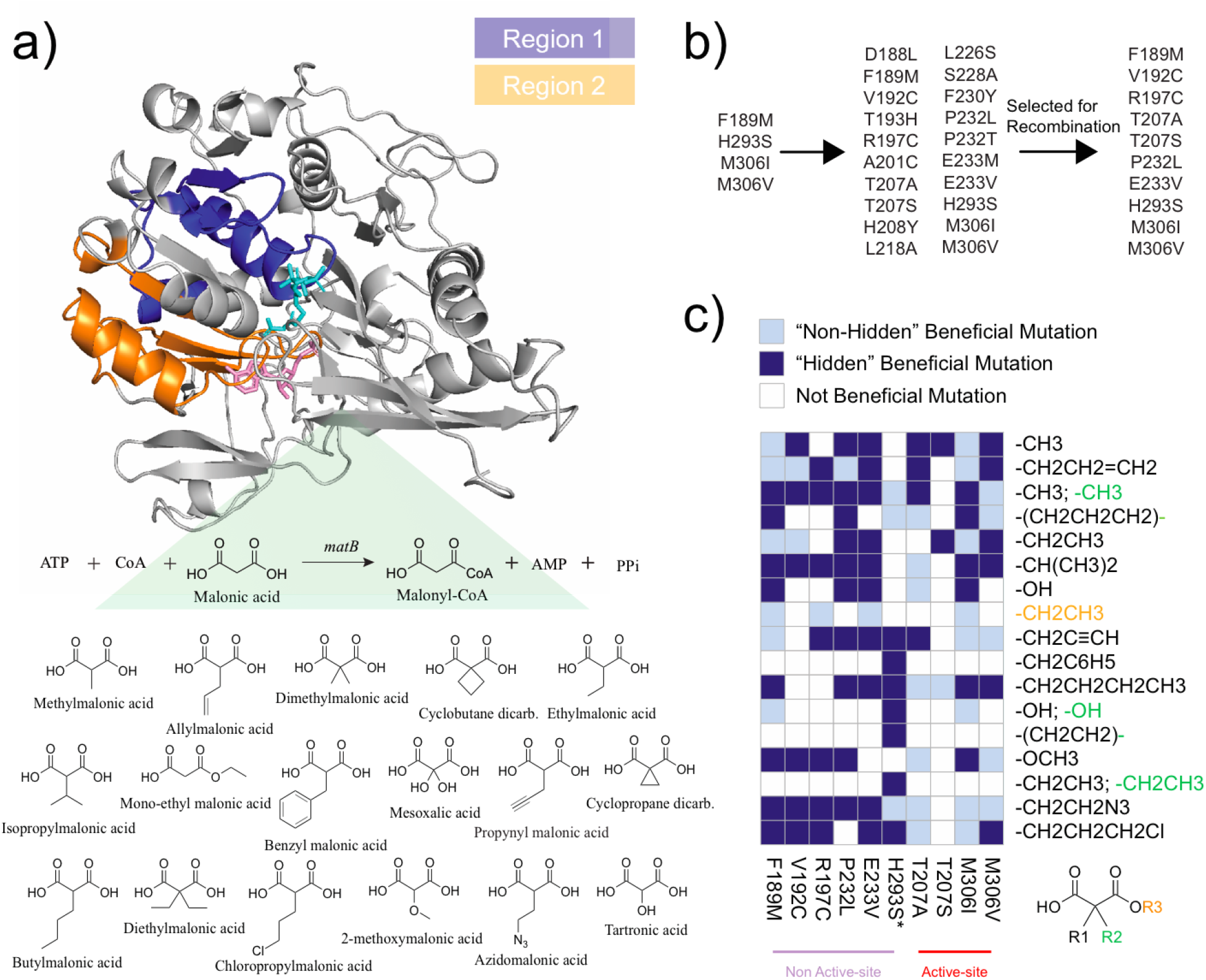
MatB screening results. **a:** Two regions flanking the active site were selected as mutagenesis targets (blue and orange), and 17 analogues of malonate were included as substrates. **b:** 20 mutations were identified in two rounds of single mutation scanning, of which 10 mutations were selected to create a recombinant library. **c:** Summary of the screening results. (1) Light blue indicates that the corresponding mutation was identified in the single mutation scanning screen for the corresponding substrate; (2) Dark blue indicates that the corresponding mutation was present in the recombinant mutants identified for the corresponding substrate, but did not appear beneficial in the single mutation scanning screen for the same corresponding substrate; (3) white if no beneficial mutation was found.

## Materials & Methods

### Reagents & Media

Unless otherwise stated, all materials and reagents were of the highest grade possible and purchased from Sigma (St. Louis, MO). Luria Broth was used for plasmid propagation and construction. Working antibiotic concentrations were as follows: spectinomycin (100 μg/mL). Components for minimum media used for protein expression were as follows: SM10++ media (pH=6.8) consists of 9 g/L (NH_4_)_2_SO_4_, 0.25 g/L citric acid, 5 mM potassium phosphate, 0.0024 g/L CoSO_4_·7H_2_O, 0.02 g/L CuSO_4_o5H_2_O, 0.0024 g/L ZnSO_4_o7H_2_O, 0.0008 g/L Na_2_MoO_4_·2H_2_O, 0.0004 g/L H_3_BO_3_, 0.0012 g/L MnSO_4_·H_2_O, 0.044 g/L FeSO_4_·7H_2_O, 2.5 mM MgSO_4_, 0.06 mM CaSO_4_, 45 g/L glucose, 200 mM MOPS (3-(N-morpholino)propanesulfonic acid), 0.01g/L Thiamine-HCl, 2.5 g/L yeast extract, 2.5 g/L casamino acids. FGM3 no phosphate Wash media consists of 3 g/L (NH_4_)_2_SO_4_, 0.15 g/L citric acid. FGM3 no phosphate Production media consists of 3 g/L (NH_4_)_2_SO_4_, 0.15 g/L citric acid, 0.0012 g/L CoSO_4_·7H_2_O, 0.01 g/L CuSO_4_·5H_2_O, 0.0012 g/L ZnSO_4_·7H_2_O, 0.0004 g/L Na_2_MoO_4_·2H_2_O, 0.0002 g/L H_3_BO_3_, 0.0006 g/L MnSO_4_·H_2_O, 0.022g/L FeSO_4_·7H_2_O, 2mM MgSO_4_, 0.05 mM CaSO_4_, 25 g/L glucose, 200 mM MOPS (3-(N-morpholino)propanesulfonic acid), 0.01g/L Thiamine-HCl. One liter of NZY^+^ media consists of: 10 g of NZ amine (casein hydrolysate), 5 g of yeast extract, 12.5 ml of 1 M MgCl2, 12.5 ml of 1 M MgSO4 and 20 ml of 20% (w/v) glucose.

### Synthesis of Malonate Analogues

2-methoxymalonic acid, 2-(pro-2yn-1-yl)malonic acid and 2-(2-azidoethyl)malonic acid were synthesized and purified following published procedures.^34^ 3-chloropropyl malonate was synthesized from Diethyl (3-chloropropyl)malonate following the procedure for 2-methoxymalonic acid, additional malonate analogues were purchased.

### Plasmid construction

Malonyl-CoA synthetase (matB) from *Rhizobium leguminosarum* was codon optimized using Codon Optimization Tool from IDT and ordered as gBlock gene fragment (IDT, Coralville, IA). MatB was cloned into pCDF under insulated low phosphate inducible promoter yibDp. pCDF-IN:yibDp-MatB has been deposited at Addgene (89631).

### Structural Modeling

Homology model for MatB from *Rhizobium trifolii* was created using SWISS-MODEL.^35^ The MatB homology model was overlaid with PDB 3NYQ.^36^ Two contiguous mutational regions (loops) each of 50 amino acids long were selected based on their proximity to the methylmalonyl-CoA substrate in the homology model for saturation mutagenesis. Specifically, Loop 1 from amino acid 184 to 234 and Loop 2 from amino acid 266 to 316, were mutated. Additionally, a mutant in Loop 1 (M306I, previously identified as a key mutant to expand MatB substrate specificity,^37^) was used as the template for Loop 2 mutations.

### Targeted Scanning Mutagenesis

Scanning mutant libraries were constructed using saturation mutagenesis using the QuikChange HT Protein Engineering System from Agilent following the manufacturer’s protocol. Mutagenic oligos were designed using eArray from Agilent. (Agilent Technologies, Santa Clara, CA). The success of the mutagenesis reaction was determined by DNA sequencing of plasmids from single colonies. For Loop 1, 100% of the sequenced clones were mutants, and 75% of them contain a single mutation. For Loop 2, 100% of the sequenced clones were mutants, and 83.3% of them contain a single mutation.

### Library Transformations

After mutagenesis, 1.5 μL of the mutagenesis reaction was transformed into SoloPack Gold supercompetent cells following the manufacturer’s protocol (Product Number 230350, Agilent, Wilmington, DE). After a 1 hour recovery, 10 μL of cells and 90 μL NZY^+^ media were plated on a prewarmed LB-Agar plate containing 100 μg/mL spectinomycin and incubated at 37 °C overnight. Colonies from all plates were counted to determine the total number of transformants per transformation, which was 14643. Transformants were diluted into NZY+ media, to obtain ~25 mutants per well in 96-well plate screens. The inoculated plate was cultured at 37 °C, 1100 rpm for 16 hours using a Mini Shaking Incubator (VWR Catalog # 12620-942, VWR International LLC., Radnor, PA, USA.). 100 μL of sterile 20% glycerol was added to 100 μL of overnight culture to prepare glycerol stock plate.

### Library Growth, Expression and Lysis

5 μL of glycerol stock was used to inoculate 150 μL SM10++ media with 100 μg/mL of spectinomycin, cultured at 37 °C, 400 rpm for 16 hours, the shaker orbit was 25 mm, using Enzyscreen plate covers (Enzyscreen, Heemstede, The netherlands). The culture was then centrifuged to remove supernatant, washed with FGM3 No Phosphate Wash, resuspended in 150 μL FGM3 No Phosphate Production media with 100 μg/mL of spectinomycin, cultured at 37 °C, 400 rpm for 24 hours for protein production. OD600 was measured for the production culture. The culture was centrifuged to remove supernatant, resuspended in 80 μL lysis buffer (100 mM Tris, pH8.0), followed by addition of 70 μL freshly prepared 4.29 mg/mL lysozyme solution (100 mM Tris, pH8.0) (final lysozyme concentration 2 mg/mL). The lysis mixture was transferred to a round bottom plate, kept at −80 °C overnight and then thawed at room temperature for one cycle the next day, centrifuged at 4000 rpm for 10 min to collect supernatant which was used in activity assays.

### Screening Activity Assays

The total volume for 96 well plate based assays was 175 μL per well, containing 50 mM sodium phosphate pH 7.2, 10 mM MgCl_2_, 0.5 mM ATP, 0.1 mM CoA, and a mixture of malonate analogues each analogue at 0.2 mM, or alternatively 0.5 mM of individual substrates. Reaction was initiated by adding 10 μL of cell lysate. The reaction was incubated at room temperature in for 4 hours, after which 25 μL of 4 mg/mL 5,5’-Dithiobis(2-nitrobenzoic acid) (DNTB) was then added, and the absorbance at 412 nm was recorded. DNTB solution was prepared fresh before use by dissolving the DNTB powder in 50 mM sodium phosphate pH 7.2. CoA consumed was calculated using a CoA standard curve. Baseline activity of wild type matB lysates was measured for the pooled substrate mixture. In the initial targeted scanning screens, mutant pools with z scores ≥3 were selected for colony isolation, evaluation with individual substrate and colony sequencing.

### Colony Isolation

Mutant pools were diluted and plated onto LB agar plates, 94 individual colonies were picked into each 96 well plate containing SM10++ with appropriate antibiotic from each agar plate, wild type MatB and pCDF empty vector controls were cultured in the remaining two wells respectively, the 96 well plates were incubated at 37 °C, 400 rpm for 16 hours. 100 μL of sterile 20% glycerol was added to 100 μL of overnight culture to prepare glycerol stock plate.

### Combinatorial Mutants

To evaluate the combined effects of identified MatB mutants, complete combinations of the mutants were prepared by first PCR amplification of sequence regions containing selected mutants using mixtures of primers and Gibson Assembly. Primers used for PCR amplification are listed in Supplemental Table S4. Due to the estimated high enzymatic activity of the combinatorial mutant pools, these pools were only screened with individual substrates. The top 5 mutant pools for each malonate analogue were selected for colony isolation and screening of individual colonies.

### Expression and Purification

To express and purify malonyl-CoA synthetase and its mutants, glycerol stock of each *E. coli* overexpressing strain was used to inoculate a flask of 50 ml SM10++ media with spectinomycin. Cells were cultured at 37°C for 16 hours. To induce the expression, the cells were collected by centrifugation, washed with FGM3 No Phosphate media, resuspended in 50 ml FGM3 No Phosphate Production media, and cultured at 30°C. After 24 hours cells were collected by centrifugation, the cell pellet was resuspended in wash buffer (20 mM phosphate, pH 7.4, 25 mM imidazole, 500 mM NaCl) and lysed by sonication. The lysate was centrifuged and the supernatant was applied to Amintra^®^ Ni-NTA resin (Expedeon, #ANN0025) and let bind for 30 min at 4°C. The resin was then washed with the wash buffer for 3 times and the protein was eluted with the elution buffer (20 mM phosphate, pH 7.4, 200 mM imidazole, 500 mM NaCl). The eluted protein was concentrated and buffer exchanged into PBS (10% glycerol) using an Amicon Ultra-0.5 3K centrifugal filters (Millipore Sigma, #C82301) and then frozen at −80°C. The purity of the protein was determined by SDS-PAGE and ImageJ analysis. The concentration of the protein was measured by Bradford assay (Thermo Scientific, #23236).

### Kinetic Analysis of Purified Enzymes

Malonyl-CoA synthetase activity was assayed with small modifications to previously reported method ^38^. Briefly, the formation of AMP in the reaction was coupled to pyruvate formation by purified myokinase and pyruvate kinase. The pyruvate formation was then coupled to NADH oxidation by lactate dehydrogenase. The oxidation of NADH was monitored at 340 nm. Assays were performed at 30°C in a total volume of 50 μl containing 100 mM HEPES, pH 7.4, 1 mM TCEP, 2.5 mM ATP, 5 mM MgCl_2_, 1 mM phosphoenolpyruvate, 0.3 mM NADH, 0.5 mM CoA, 0.125 U myokinase, 0.9 U pyruvate kinase, 0.65 U lactate dehydrogenase, malonate or the corresponding malonate analog (starting with 25 μM - 1 mM and repeating with 1 mM - 16 mM if not saturating). The reaction was initiated with the addition of MatB or the corresponding mutant (starting with 25 nM and repeating with 500 nM if no activity detected at lower enzyme levels). Reactions were monitored at 340 nm for 1 hour (30s per read) using a SpectraMax Plus 384 microplate reader (Molecular Devices). NADH concentrations were determined by its extinction coefficients. ^39^ Linear regressions were performed on a sliding window with a width of 10 data points scanning the change of NADH concentration over time. The initial velocity of each reaction was obtained as the most negative slope estimation among its regression models that have a R^2^ greater than 0.9. The Vmax and Km of malonyl-CoA each synthetase and malonate analogue was determined by fitting the data to the Eadie-Hofstee equation (v = −Km*v/[S] + Vmax, where v is the initial velocity and [S] is the substrate concentration) using the ordinary least square estimator in statsmodels, a Python package. The kcat of each pair was calculated as Vmax/[E], where [E] is enzyme concentration in the reaction. The error of kcat was determined by error propagation of errors from Vmax and [E]. Data are reported as mean ± s.e. (n = 3). P-values were determined by student t tests.

### Structural Modeling of Mutants

Structural modeling of mutations was initially performed on KING^40^ and PyMOL^41^. Cavity mapping was performed by the FOO function of MAGE^42^. Determination of atomic contact between residues was achieved by Arpeggio.^43^ Visualization of atomic contacts and prediction of free energy change of mutations were carried out by PremPS. ^44^

## Results

We chose malonyl-CoA synthetase from *Rhizobium trifolii* as our focus of this study for three primary reasons: firstly, it has some natural substrate promiscuity, ^36^ making it a good starting point for the directed evolution of substrate specificity. ^45,46^ Secondly, it catalyzes the formation of malonyl-CoA, a very important precursor in polyketide synthesis. ^36^ The engineering of enzyme variants capable of the activation of diverse malonate analogues has the potential to enable the biosynthesis of diversified natural product analogues.^33^ Thirdly, this synthetase activity can be rapidly evaluated using a high-throughput colorimetric screening assay, based on quantification of the CoA thioester. ^47^

We purchased or synthesized a panel of 17 malonate analogues (18 substrates including malonate itself) in our study (Figure 2a). These analogues represent a diverse profile of chemical properties (i.e. esters, halogens, hydrophobic R groups, conjugation handles, etc.). We next developed a screening assay and measured the activity of the wild type malonyl-CoA synthetase from *Rhizobium trifolii* (wild type MatB) towards all selected substrates. As previously demonstrated and illustrated in Figure S2, wild type MatB exhibits activity with malonate, and some activity with methylmalonate, allylmalonate, and mono-ethyl malonate, but had no measurable activity to other malonate analogues in our panel.^34^

We next generated a homology model of the wild type MatB protein. Using this model we chose two 50 amino acid long regions (region 1 and region 2, Figure 2a), which flank the active site, as regions for mutagenesis. Two scanning mutant libraries covering all possible single mutations of the two selected regions were created and screened (~1000 unique variants per region, 2000 in total). To efficiently screen multiple malonate analogues with the mutant library in parallel, we developed a pooling method in the format of 96-well plates that reduces the number of assays by over 10-fold (Figure S3). Briefly, mutants were pooled together with one well containing ~25 mutants and screened with a mix of the substrate analogues. The hit mutant pools exhibiting improved activity based on Z scores were next assayed against each individual substrate to identify the substrate(s) for which they have improved activity. Finally, mutants in each hit mutant pool were separated by colony isolation, and again assayed against the substrate(s) for which the pool improved activity, to identify single mutants. Initial screening of the two libraries led to the identification of four mutations: F189M, H293S, M306V, and M306I. Notably, M306I and M306V have been previously reported as key mutations that can alter the specificity of MatB, ^34,37^ validating that this screen was effective to identify beneficial mutations.

To identify additional beneficial mutations, we created a new scanning mutant library with the M306I variant as a template. As M306I is a mutant in region 2, the new scanning mutant library (which utilized the same mutagenic oligo pools) had all possible single mutations in region 1. We applied the same screening method to this new library and identified 16 more beneficial mutations (Figure 2a). Of these 16 new mutations, at most only 3 would have been identified when screening with individual substrates. Among these mutations, T207A and T207S were previously reported to alter the specificity of this enzyme, which once again confirms the efficacy of our screen.^34^ Both T207 and M306 mutations are active site mutations(Figure 2c), making 4 of the total 20 mutations accessible by more traditional structure-guided evolution strategies. In comparison, the other 16 mutations at 14 positions are located away from the active site, with 44% of them (7 out of 16) residing on the protein surface (Table S1). Such mutations would not be identified with structure-guided approaches that focus on residues around the active site, as previously reported.^34^ This is also consistent with a previous study stating that substrate specificity is globally encoded.^48^ Refer to Supplemental Table S2 for a summary of the evolving fitness of the mutant pools during this directed evolution.

We recombined 10 of the 20 mutations at 8 positions we identified for all substrates (Figure 2a-b), giving us a library of 576 unique recombinants. The 10 mutants were chosen based on their occurrences among substrates as well as the ease of constructing recombinant variants (Table S2). We then screened this new combinatorial library against each individual substrate and identified a total of 26 unique recombinants with improved activity for one or more of the 18 substrates. Kinetic parameters for the best performing recombinants (kcat and Km) were determined for purified enzymes and are given in Figure 3 and Table S3. As summarized in Figure 2c, the best recombinant identified to date for each substrate almost always contained mutations identified in earlier rounds due to activity with a different substrate. Such “hidden” mutations would have been missed upon screening for a single substrate alone. Take tartronic acid as an example, the recombinant mutant sequences identified for this substrate included F189M, T207A, P232L, E233V, and M306I. However, by screening single mutants of wild type MatB with tartronic acid as the only substrate (single agent), no single mutations would be identified (Figure 4). Only by screening single mutants of MatB-M306I, which is identified by screening other substrates, T207A was identified as a beneficial mutation for tartronic acid with limited improvement. Without other substrates, only double mutant screening could lead to the discovery of T207A-M306I. Nevertheless, to identify these recombinant sequences, iterative rounds of mutagenesis and screening would have been needed to identify F189M, P232L, and E233V (Figure 4a). In other words, the recombinants we identified for tartronic acid would be extremely difficult if not impossible to identify using the traditional iterative single mutation scanning or a site-saturation mutagenesis strategy generating large combinatorial libraries. By incorporating substrate diversity (multiple agents) into the screening, this approach enables the identification of a broader set of mutations that have a higher likelihood to impact enzyme activity when in combination with other mutations, leading to a prioritization of the combinatorial space and more rapid identification of the combinatorial mutants. Moreover, 20 of the 26 single mutations used in the recombinant pool were only identified for one substrate (Table S2), suggesting that we did not just screen for more promiscuous variants that have broadly improved activity with multiple substrates.

**Figure 3.**
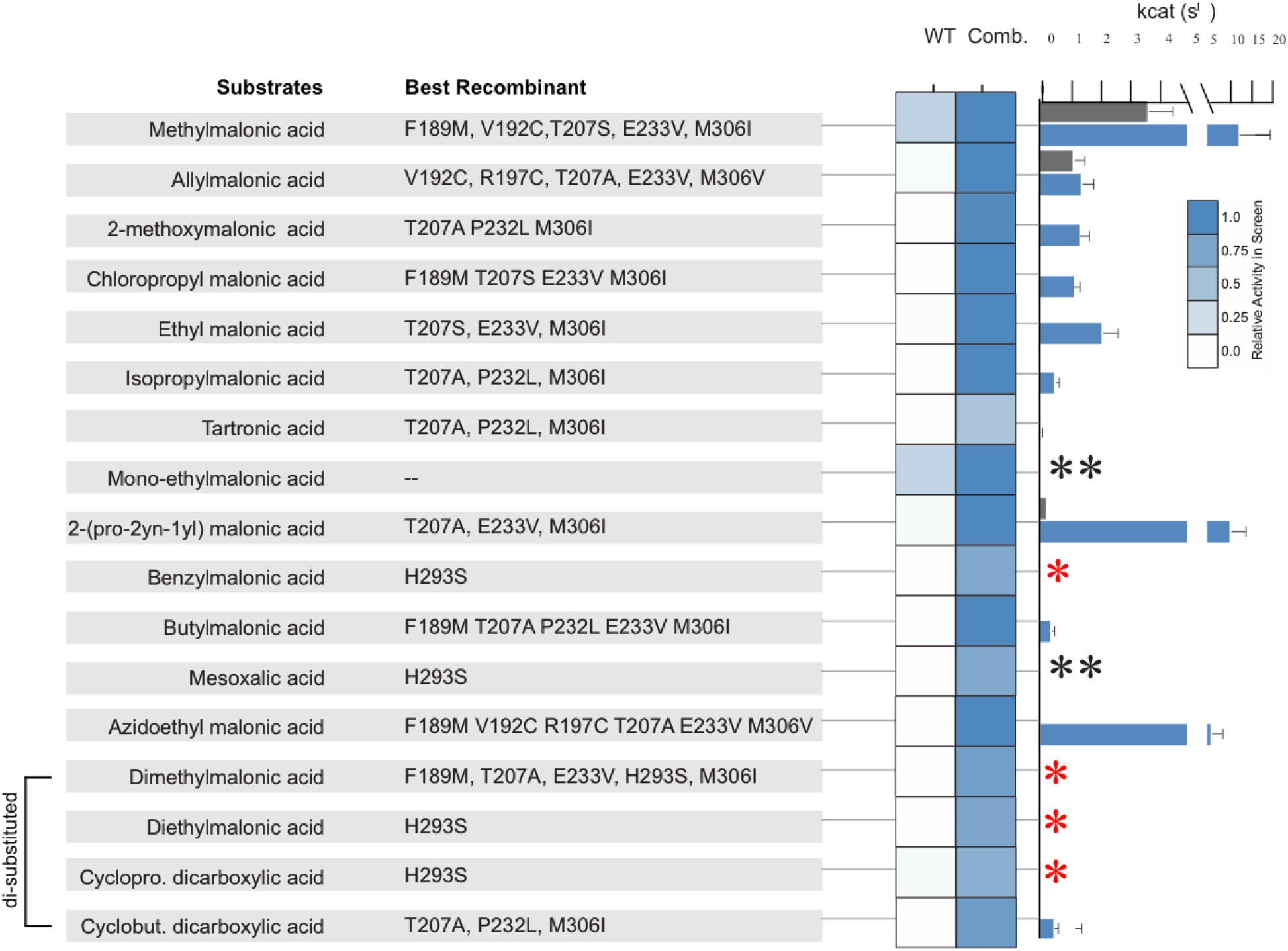
Kinetic characterization of purified mutant enzymes. The colormap indicates the activity of the substrate-recombinant pair determined in the screening assay, while the bar chart indicates the kcat of substrate-recombinant pair determined in confirmatory purified enzyme assays. A red asterisk indicates that activity not detected in the confirmatory enzyme assay. A double black asterisk the enzyme assay was not successful due to (1) substrate hydrolysis of mono-ethylmalonic acid; (2) reaction between substrate and a coupling enzyme in the assay for mesoxalic acid (Figure S4).

**Figure 4.**
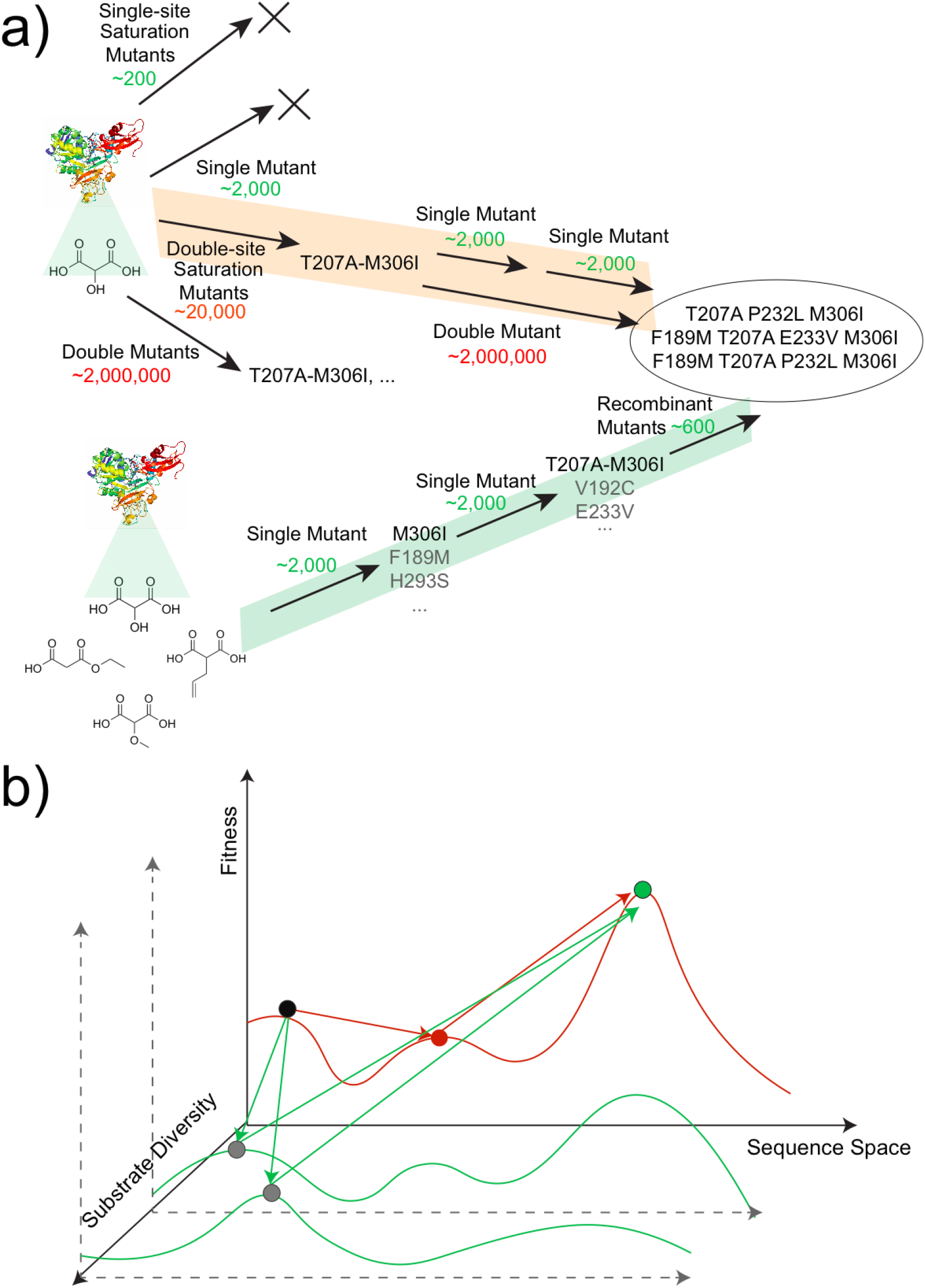
Evolutionary routes from wild type MatB to the recombinants identified with improved activity for tartronic acid. **a:** In the single agent setting (orange path), single-site saturation and single mutant scanning strategies are comfortably tractable (green numbers), but will not lead to the discovery of beneficial mutations that were identified. Double-site saturation mutagenesis focused on the active site residues is still manageable and will lead to identification of the double mutant T207A-M306I, but it will miss mutations at other sites. To identify the best recombinant mutants, additional rounds of mutation and luck to avoid any fitness valley are needed (the orange route) on top of double-site saturation screening. The complete evaluation of double mutants is less tractable (red numbers). In comparison, in the multi-agent setting (green path), screening of the single mutant library of moderate size will lead to a set of mutations potentially impacting the activity on tartronic acid. One single step of screening of a small library created by combining the mutations will lead to the identification of the recombinants conveniently. **b:** Additional agents can provide more efficient routes to optimal solutions (green circle) by sampling additional dimensions of data (green lines). Using mutants identified with multiple agents (gray circle) it is possible to “hop”/ bypass potential fitness valleys that may exist for a single agent (orange line).

We next sought to better understand how these mutations may be impacting the activity of MatB. We can categorize the mutations into two classes: (i) active-site mutations, including mutations at positions T207 and M306, and (ii) non-active-site mutations, including all other mutations, and importantly most of the hidden mutations (Figure 2c). We analyzed the expected structural impact of T207A and M306I mutations in our homology model. These mutations allow more space for the C-2 substituted carbon of malonyl-CoA, thus enabling the activation of malonate analogues with larger side chains (Figure 5a). The binding pocket is enlarged by mutating the relatively bulkier threonine and methionine to smaller alanine and isoleucine, respectively (Figure S5).

**Figure 5.**
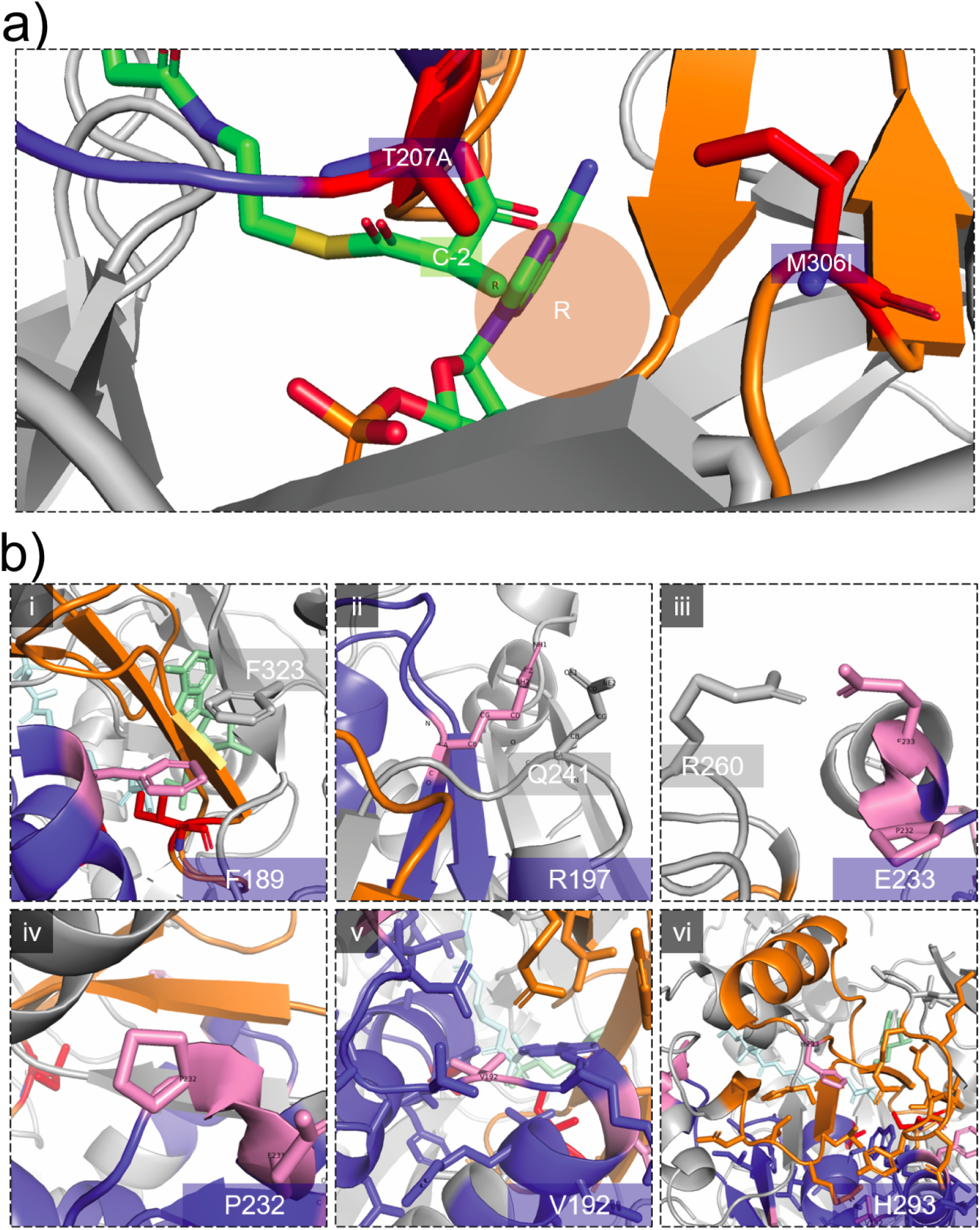
An overview of proposed mechanisms underlying the impact of key mutations. **a:** T207A and M306I (in red) are mutations that are expected to create more space for the C-2 side chain. **b:** Mutants at positions (i) F189, (ii) R197, and (iii) E233 are expected to form side chain interaction with remote residues (gray), which would be potentially interrupted by a mutation. (iv) Residue P232 is a surface proline residue that imposes restriction on the enzyme backbone that would be alleviated by a mutation. (v) Residue V192 is expected to form hydrophobic interactions with a group of mostly hydrophobic side chains surrounding it. (vi) Residue H293 is also surrounded by a group of residues with various properties.

As for the non-active site mutations, we picked a few representative ones for our analysis, including F189M, V192C, R197C, P232L, P232T, E233V, and E233M. Initially, we analyzed these mutations in our homology model. As illustrated in Figure 5b, we found that these residues are all involved in some kind of side chain interactions, except P232. Specifically, F189 is closely located to another surface residue, F323(Figure 5b(i)). These two phenylalanines would be likely to form a π-π interaction between two alpha helices. We reasoned that this interaction would be disrupted with the F189M mutation, presumably destabilizing the interaction between these two helices. Similarly, R197 resides closely with Q241, and E233V resides closely with R260 (Figure 5b(ii)& 5b(iii)). These two residues are likely to be engaged in stabilizing interactions between secondary structural elements through hydrogen bonding, which again would be disrupted when mutated. V192 is seated in a cluster of hydrophobic side chains(Figure 5b(v)), where we reasoned that a mutation to cysteine would be greatly unfavored by the surrounding residues.

Motivated by these observations, as shown in Figure 6, we computationally predicted the atomic contacts of these residues in both wild-type and mutant sequences. We discriminated between local (residues within 6 amino acids on the primary sequence) and remote contacts. Additionally, we predicted the free energy change caused by these mutations (Table S1). Among the 5 mutations subject to this analysis (excluding P232L and P232T), all mutations display a reduced number of remote contacts and thus total contacts formed by the residue (Figure 6d), with 4 of them exhibiting increased free energy (Table S1). This suggests a loss of remote stabilizing interactions. In contrast, P232 is a unique surface residue that imposes a strict restriction of the conformation of enzyme backbone due to the unique ring structure of proline. Mutation of the proline is expected to increase backbone flexibility. We hypothesize that these residues represent stabilizing surface interactions limiting conformational changes. When such interactions are disrupted by mutations, the enzyme is likely to become more flexible, thus enabling altered specificity. These impacts are magnified when paired with other mutations. Better validation of this mechanism requires further study. Additionally, while the mutation H293S was identified in initial screens and included in subsequent libraries, we were never able to confirm an improvement in enzyme activity in our kinetic assays of purified variants. As a result, we currently consider this mutation a false positive in our screen, where further effort is needed to elucidate its impact/observed advantage in the screening assay.

**Figure 6.**
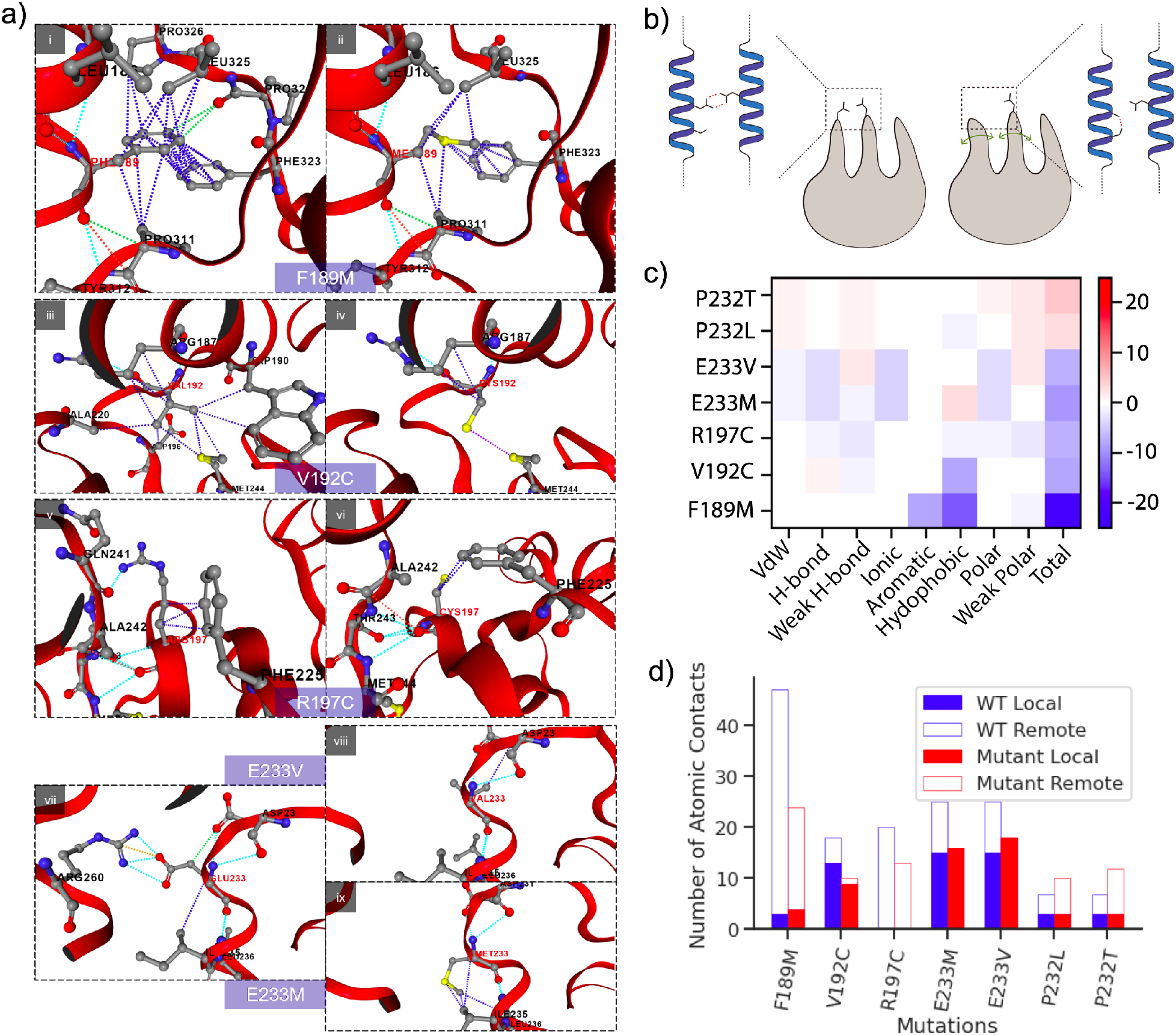
A structural analysis of mutations. **a:** Visualization of atomic contacts formed by the wild-type and mutant residue of each beneficial mutation. (i)&(ii): hydrophobic and aromatic interactions formed by F189 and F323 are largely disrupted by F189M; (iii)&(iv): hydrophobic interactions formed by V192 and its surrounding residues are reduced with a V192C mutation; (v)&(vi): The number of interactions formed by R197 and F225 are reduced by R197C, and the interaction with the carbonyl oxygen of Q241 is eliminated in the mutant; (vii)-(ix): The interactions between E233 and R260 are eliminated in both E233V and E233M. **b:** A schematic representation of how the loss of remote contacts could contribute to enhanced enzyme flexibility. **c:** The change in numbers of atomic contacts caused by the mutations differentiated by the type of interaction. **d:** The change in numbers of atomic contacts caused by the mutations differentiated by range of interactions (local vs. remote). The numbers of total contacts and remote contacts are reduced by all mutations except P232L and P233T.

Another interesting result we observed is that virtually no improved mutants were identified for most di-substituted substrates (Figure 3). When examining the structural model, and the enzyme complex with ATP/AMP, we hypothesize that it might be intrinsically challenging for this enzyme to bind with both C-2 di-substituted malonates as well as ATP as the addition of a second R-group to the C-2 position of malonyl-CoA, is expected to produce a steric clash with ATP/AMP (Figure S6). This may also provide a mechanism for this enzyme to manage its enantioselectivity.

## Discussion

We have demonstrated improved efficiency in directed evolution by leveraging multi-agent searching, in this case substrate diversity. By interrogating MatB mutant libraries of modest size (~1000 variants per library) with a panel of diverse malonate analogues, we successfully obtained a set of 20 mutations altering substrate specificity, which is much larger than the set of beneficial mutations we would have obtained by screening only one substrate (a maximum of 4 in these results). Subsequently, screening a relatively small recombinant library of only 576 unique variants led to the identification of improved recombinants for many of the substrates. Using this multi-agent approach, we are able to extract additional information without aggressively expanding the scope of genetic diversity being screened. This is particularly important because with many current screening methods even the complete evaluation of the double mutant space can be experimentally intractable if the targeted region of mutagenesis is relatively large (Figure S1). The additional information from alternative substrates can lead to the discovery of more potentially beneficial mutations in fewer rounds without sacrificing the number of residues in the scope of the search. More importantly, the additional information provides alternative routes on the fitness landscape that could bypass valleys in any given trajectory for a single substrate (Figures 1 and 4b). In addition, although the approach can increase the initial number of screening assays, even with the pooling method we developed, it still saves a significant amount of time and effort by reducing the need of library cloning as fewer rounds of screening are needed. We expect this approach to complement other strategies which focus on using structural knowledge or statistical analysis to prioritize the sequence space^4,28^ or alternatively on expanding the throughput.^20^ To our knowledge, this is the first time that substrate diversity has been strategically incorporated in directed evolution and that multi-agent searching has been shown to identify hidden mutations.

Additionally, we were somewhat surprised by the discovery that a majority of the identified beneficial hidden mutations reside far from the active site on the surface of MatB. While the stability of tertiary or quaternary structures are often engineered for improved protein expression, solubility or stability, ^49,50^ to our knowledge reducing stability as a route to enhance substrate specificity has not been previously appreciated. This is despite the fact that these types of mutations are more commonly identified in screens leveraging random mutagenesis.^28^ In the case of MatB, we hypothesize that these hidden mutations selectively destabilize interactions between elements of secondary and/or tertiary structure, implicating the potential that changes in protein flexibility may be of equal importance to changes of more fixed structural elements of the active site. This is an important hypothesis requiring significant further effort, as these types of changes may well be predictable with future algorithms. If so these insights may enable additional structure guided approaches focusing on areas of structure outside of the active site.

Despite the success of this work, there are limitations of the approach as implemented to date. Firstly, like many other screens, the presence of false positives is of concern, in our case the H293S mutant whose impact on activity could not be confirmed in subsequent assays. Secondly, no improvements in activity were found for 4 substrates in our panel. Notably, 3 of them are C-2 disubstituted, and the other has a bulky benzyl group on the side chain. These structures may represent diversity too remote from the native substrate structure to be useful in the approach. Obviously, the selection of the substrate analogue panel is crucial for the effectiveness of the approach. This raises a future challenge in defining “criteria” for the selection of good substrate analogues. In this study, we selected our substrate panel by their accessibility (commercial availability and/or simplicity of synthesis). In the future, a better selection criteria integrating quantification of the similarity to the native structure should be investigated and applied to this approach to improve it.

Lastly, the use of a multi-agent searching has the potential to improve the efficiency of searching sparse fitness landscapes in the engineering of other enzymes and proteins. Sparse fitness landscapes are also more common in biological systems and a challenge in other bioengineering efforts, from synthetic biology to metabolic engineering. Generically, the use of multi-agent searching, beyond the case of multiple substrates, may enable improved methods for finding combinatorial optimum.

## Acknowledgements

We would like to acknowledge the following support: DARPA# HR0011-14-C-0075, ONR YIP #N00014-16-1-2558, and DOE EERE grant #EE0007563. Additionally we would like to thank Dr. Jane S. Richardson and Dr. David C. Richardson for their support in structural modeling.

## Author contributions

T. Yang, Z. Ye, and M.D. Lynch designed and analyzed experiments, and performed analytical analyses. T. Yang and Z. Ye constructed the mutant libraries and performed activity screenings. T. Yang ran computational analyses of mutants. All authors wrote, revised and edited the manuscript.

