## Supplementary Materials for "“Multi-Agent” Screening Improves the Efficiency of Directed Enzyme Evolution"

### Supplemental Materials

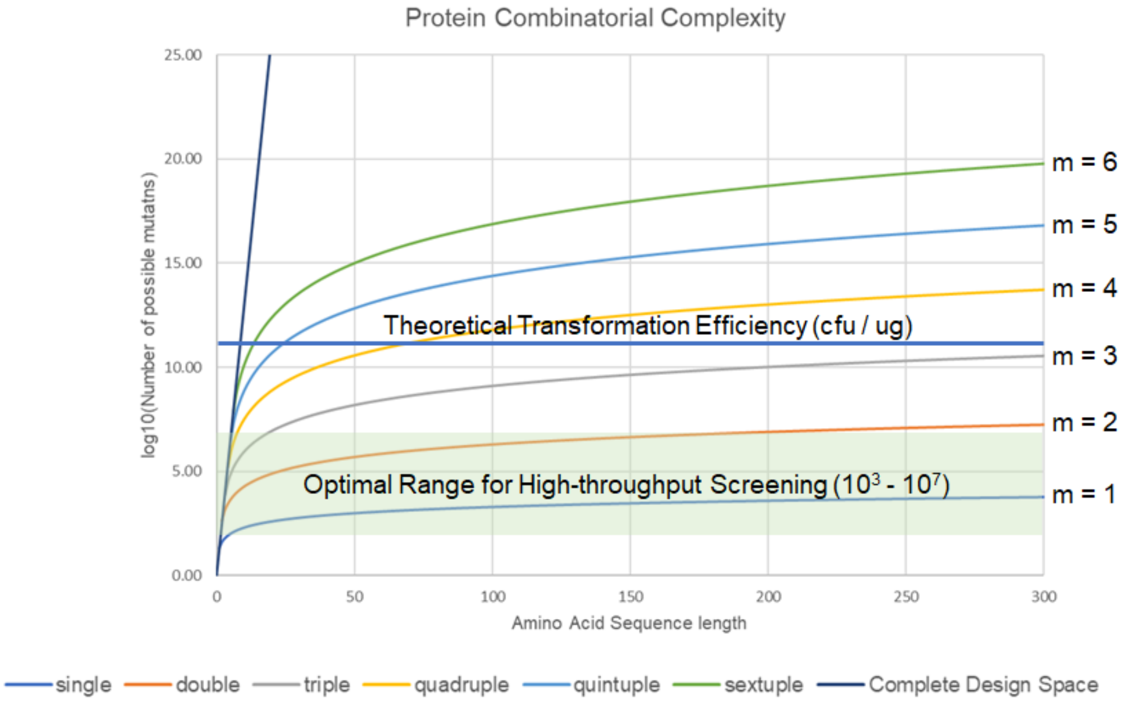


**Figure S1:** The Complexity of Protein Design Space with Different Criteria. The complexity of complete design space grows rapidly out of the range for high-throughput screening at very small sequence length. The curves designated with an m value represent the complexity of the m-mutant space at different sequence length, with only single- and double-mutant space manageable for the high-throughput screening capacity. The theoretical transformation efficiency is also labeled in the plot for reference.

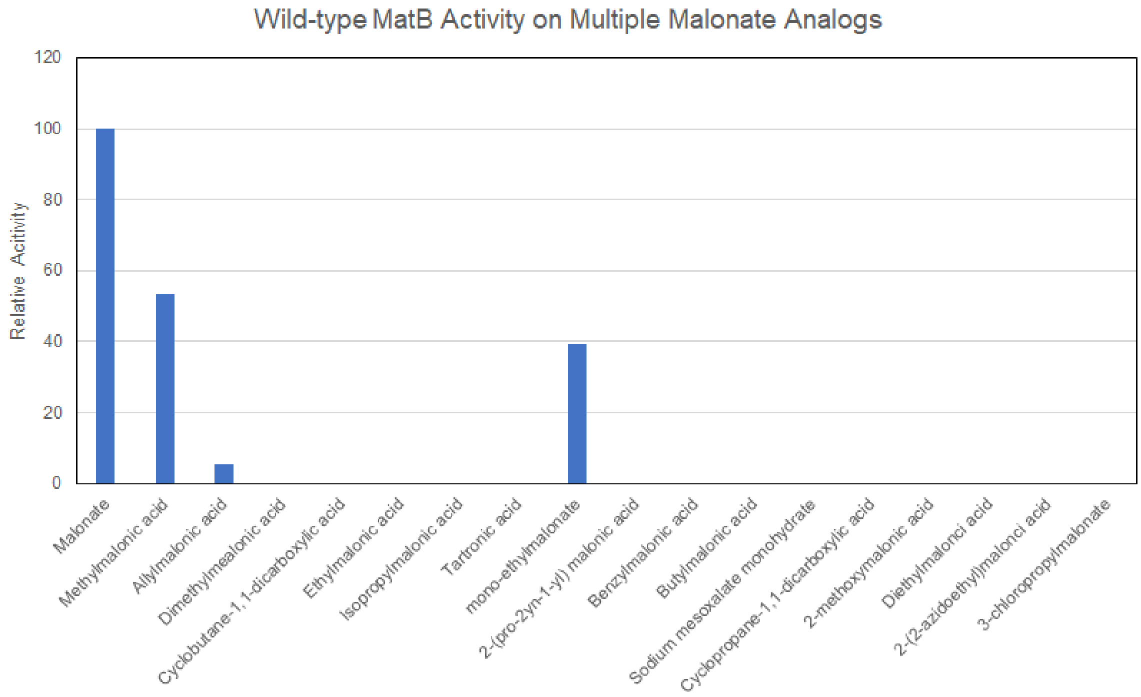


**Figure S2.** WT MatB Screening Activity Profile on the Selected Panel of Substrate Analogues.

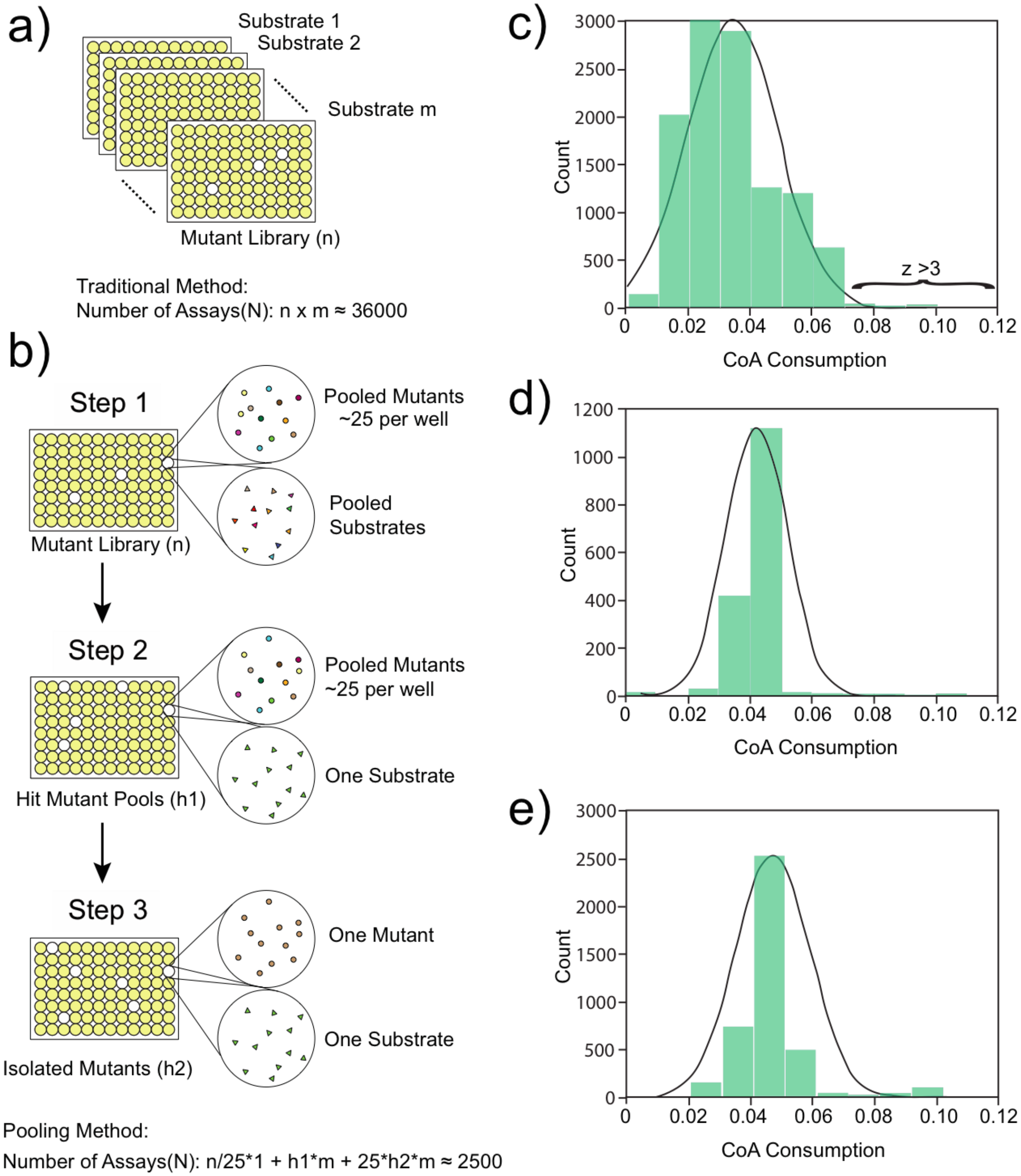


**Figure S3. A comparison of screening methods and results for MatB. a:** a "naive" method to screen multiple substrates in parallel, where number of assays (N) scale linearly with number of substrates (m) and library size (n). **b:** A more efficient pooling method. In the first step, mutants were pooled together, with one well containing 25 mutants. We assay the mutant pools against the mix of the substrate analogs. After selecting the hit mutant pools that show improved activity on the substrate mix, we assay the hit mutant pools against each individual substrate in the second step to find out which substrate each mutant pool has improved activity for. Then, we separate the mutants in each hit mutant pool by colony isolation, and we assay individual mutants in each pool against the substrates that the pool shows improved activity on to identify the single hit mutants of each substrate. The pooling method reduces the number of assays by more than 10-fold. **c-e:** Distribution of activity readout in each step (c: step 1, d: step 2, e: step 3) of the pooling screen for MatB. The assayed population is gradually moving toward higher activity.

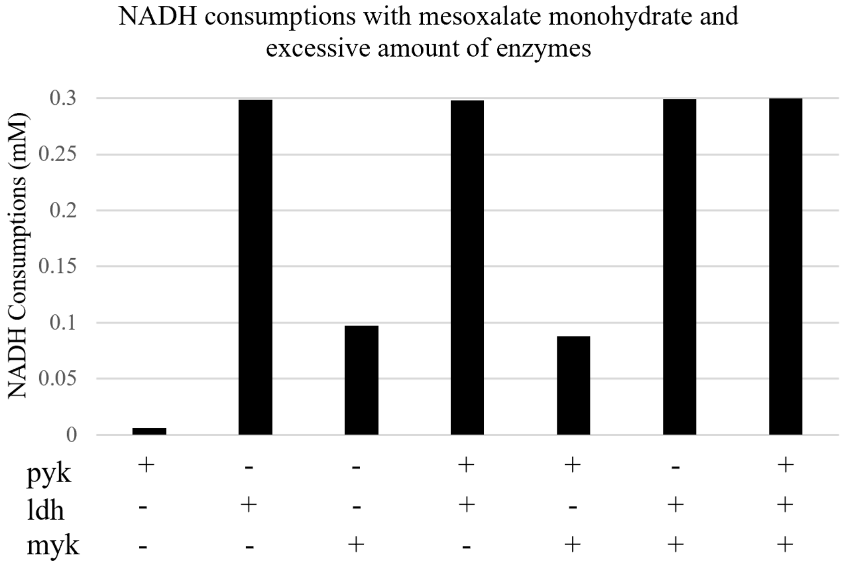


**Figure S4:** Reaction of Mesoxalic acid with NADH in the presence of coupling enzymes. The formation of AMP in the synthetase (matB) reaction is coupled to NADH oxidation by purified myokinase, pyruvate kinase, and lactate dehydrogenase, which can be monitored at 340 nm. Abbr.: pyk (pyruvate kinase), ldh (lactate dehydrogenase), myk (myokinase).

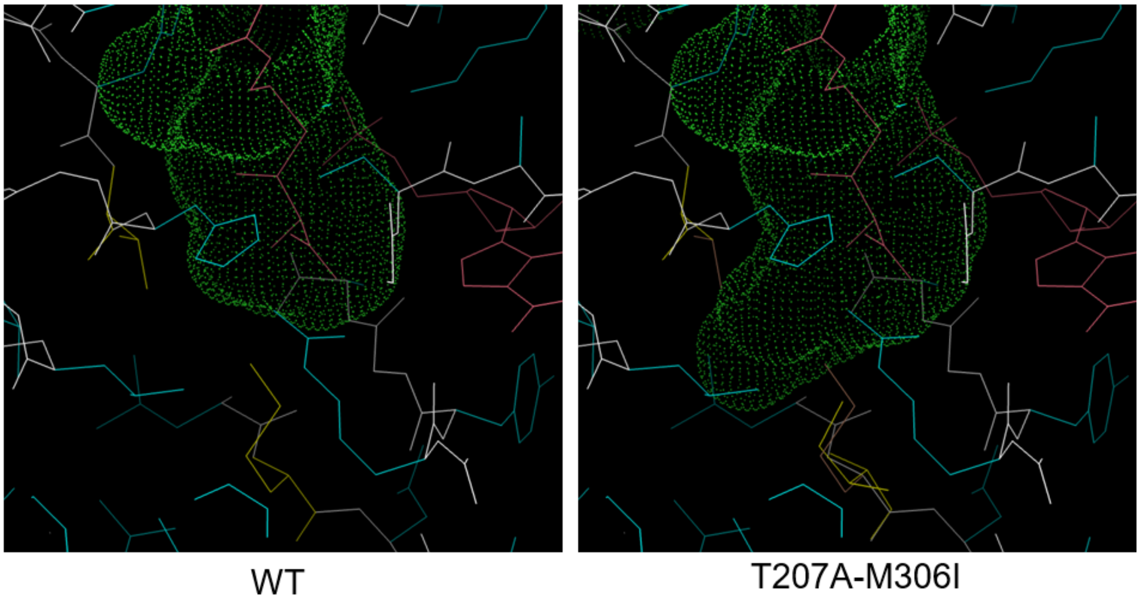


**Figure S5** Enlarged Cavity of MatB Binding Pocket(green dots) of T207A-M306I.

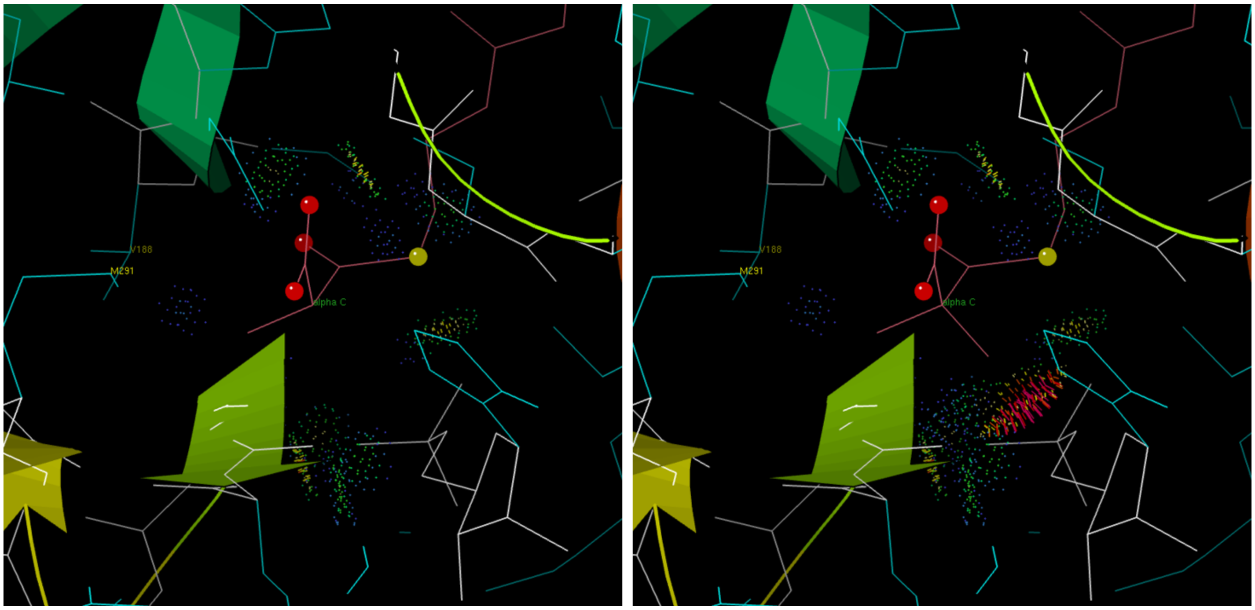

**Figure S6.** Clash between the introduced second substitution group and the AMP molecule in the homology model.

| **Table S1. Predicted ΔΔG and locations of mutations** | | |
|---|---|---|
| **Mutation** | **Predicted ΔΔG/ kcal·mol^-1** | **Location** |
| D188L | 0.44 | SUR |
| F189M | 1.04 | COR |
| V192C | 1.3 | COR |
| T193H | 1.01 | SUR |
| R197C | -0.23 | SUR |
| A201C | 0.8 | COR |
| H208Y | 0.23 | COR |
| L218A | 2.06 | COR |
| L226S | 2.44 | COR |
| S228A | 0.1 | SUR |
| F230Y | 1.35 | COR |
| P232L | -0.35 | COR |
| P232T | 0.65 | COR |
| E233M | 0.62 | SUR |
| E233V | 0.56 | SUR |
| H293S | 1.03 | SUR |

SUR: surface; COR: core;

| **Table S2: Summary of Mutant Progression.** | | | | | | | |
|---|---|---|---|---|---|---|---|
| **Substrate** | **Targeted Scanning Mutants** | | | | | **Mut. for Comb.** | **Combinatorial Mutants (26 unique hits)** |
| Methylmalonic acid | 4 | F189M | F230Y | P232T | L218A | P232T<br>F230Y<br>**P232L**<br>**V192C**<br>**T207A**<br>S228A<br>D188L<br>**E233V**<br>**R197C**<br>A201C<br>**E233M**<br>H208Y<br>L226S<br>T207S<br>T193H<br>**F189M** | T207A E233V M306I<br>F189M T207A P232L E233V M306V<br>F189M V192C T207S E233V M306I |
| Allylmalonic acid | 3 | M306I | V192C | P232L | | | V192C R197C T207A E233V M306<br>F189M T207A E233V M306I |
| Dimethylmalonic acid | 3 | M306V | H293S | F189M | | | F189M T207A E233V H293S M306I<br>F189M V192C R197C T207A E233V H293S M306V<br>F189M V192C R197C T207A P232L E233V M306I |
| Cyclobutane-1,1-dicarboxylic acid | 4 | M306V | H293S | F189M | T207A | | H293S<br>F189M T207A P232L M306I<br>T207A P232L M306I |
| Ethylmalonic acid | 4 | M306I | D188L | V192C | S228A | | T207S E233V M306I<br>T207A E233V M306I<br>F189M T207A P232L M306V<br>T207A P232L E233V M306I |
| Isopropylmalonic acid | 1 | T207A | | | | | T207A P232L M306I<br>F189M V192C R197C T207A P232L M306V |
| Tartronic acid | 2 | F189M | T207A | | | | T207A P232L M306I<br>F189M T207A E233V M306I<br>F189M T207A P232L M306I |
| mono-Ethyl malonate | 3 | F189M | E233V | R197C | | | |
| 2-(pro-2yn-1-yl)malonic acid | 4 | M306V | F189M | A201C | E223M | | F189M T207A P232L M306I<br>T207A E233V M306I<br>F189M R197C T207A P232L H293S M306V<br>F189M T207A E233V M306I |
| Benzylmalonic acid | 2 | H208Y | L226S | | | | H293S |
| Butylmalonic acid | 1 | T207S | | | | | T207A E233V M306I<br>T207S P232L H293S M306V<br>F189M T207A P232L E233V M306I<br>F189M T207A E233V M306V |
| Sodium mesoxalate monohydrate | 2 | F189M | T193H | | | | H293S |
| Cyclopropane-1,1-dicarboxylic acid | 0 | | | | | | H293S |
| 2-methoxymalonic acid | 3 | M306V | F189M | T207A | | | T207A P232L M306I<br>V192C T207A P232L M306I<br>F189M V192C R197C T207A P232L M306I |
| Diethylmalonic acid | 0 | | | | | | H293S |
| 2-(2-azidoethyl)malonic acid | 4 | M306V | H293S | F189M | T207A | | T207A E233V M306I<br>F189M T207A P232L E233V M306V<br>V192C T207A P232L M306I<br>F189M V192C R197C T207A E233V M306V |

| **Table S3: Kinetic Parameters of Purified Enzymes** | | | | | | |
|---|---|---|---|---|---|---|
| **Malonate Analogue** | **Best Mutant** | **Notes** | **Mutant Kcat (s^-1)** | **Mutant Km (µM)** | **Wt kcat (s^-1)** | **P Value** |
| Malonate | wt | | | | 35.8±7.7 | -- |
| Methylmalonic acid | F189M V192C T207S E233V M306I | | 12.9±7.1 | 4298±1766 | 3.72±0.82 | 0.13 |
| Allylmalonic acid | V192C R197C T207A E233V M306V | | 1.46±0.38 | 94±15 | 1.18±0.38 | 0.31 |
| Dimethylmalonic acid | F189M V192C R197C T207A E233V H293S M306V | False Positive | | | | |
| Cyclobutane-1,1-dicarboxylic acid | T207A P232L M306I | | 0.52±0.14 | 1544±437 | -- | 0.003 |
| Ethylmalonic acid | T207S E233V M306I | | 2.16±0.52 | 152±31 | -- | 0.002 |
| Isopropylmalonic acid | T207A P232L M306I | | 0.55±0.12 | 1161±188 | -- | 0.0008 |
| Tartronic acid | T207A P232L M306I | | 0.078±0.020 | 1733±435 | -- | 0.03 |
| mono-Ethyl malonate | -- | Hydrolysis of Substrate | | | | |
| 2-(propyn-1-yl)malonic acid | T207A E233V M306I | | 10.8±3.5 | 797±37 | 0.29±0.16 | 0.01 |
| Benzylmalonic acid | H293S | False Positive | | | | |
| Butylmalonic acid | F189M T207A P232L E233V M306I | | 0.42±0.09 | 139±19 | -- | 0.002 |
| Mesoxalic acid | H293S | Substrate reacts with coupling enzymes | | | | |
| Cyclopropane-1,1-dicarboxylic acid | H293S | False Positive | | | | |
| 2-methoxymalonic acid | T207A P232L M306I | | 1.42±0.32 | 1861±325 | -- | 0.002 |
| Diethylmalonic acid | H293S | False Positive | | | | |
| 2-(2-azidoethyl)malonic acid | F189M V192C R197C T207A E233V M306V | | 6.0±2.5 | 1994±789 | -- | 0.03 |
| 3-chloropropyl malonate | F189M T207S E233V M306I | | 1.2±0.3 | 211±24 | -- | 0.002 |

| **Table S4: List of primers used for selected MatB mutants combination.** | | |
|---|---|---|
| PCR # | Primer Name | Sequence |
| 1 | MatB-F189M-FOR | GCTTACATTGCGTGATATGTGGCGTGTAACCGCAGGCGATCGTCT GATTCATGCGTTACCGATTTTTCATACACATGGCC |
| | MatB-V192C-FOR | GCTTACATTGCGTGATTTCTGGCGTTGCACCGCAGGCGATCGTCT GATTCATGCGTTACCGATTTTTCATACACATGGCC |
| | MatB-R197C-FOR | GCTTACATTGCGTGATTTCTGGCGTGTAACCGCAGGCGATTGCCT GATTCATGCGTTACCGATTTTTCATACACATGGCC |
| | MatB-T207A-FOR | GCTTACATTGCGTGATTTCTGGCGTGTAACCGCAGGCGATCGTCT GATTCATGCGTTACCGATTTTTCATGCGCATGGCC |
| | MatB-F189M-V192C-FOR | GCTTACATTGCGTGATATGTGGCGTTGCACCGCAGGCGATCGTCT GATTCATGCGTTACCGATTTTTCATACACATGGCC |
| | MatB-F189M-R197C-FOR | GCTTACATTGCGTGATATGTGGCGTGTAACCGCAGGCGATTGCCT GATTCATGCGTTACCGATTTTTCATACACATGGCC |

| | | |
|---|---|---|
| | MatB-F189M-T207A-FOR | GCTTACATTGCGTGATATGTGGCGTGTAACCGCAGGCGATCGTCTGATTCATGCGTTACCGATTTTTCATGCGCATGGCC |
| | MatB-V192C-R197C-FOR | GCTTACATTGCGTGATTTCTGGCGTTGCACCGCAGGCGATTGCCTGATTCATGCGTTACCGATTTTTCATACACATGGCC |
| | MatB-V192C-T207A-FOR | GCTTACATTGCGTGATTTCTGGCGTTGCACCGCAGGCGATCGTCTGATTCATGCGTTACCGATTTTTCATGCGCATGGCC |
| | MatB-R197C-T207A-FOR | GCTTACATTGCGTGATTTCTGGCGTGTAACCGCAGGCGATTGCCTGATTCATGCGTTACCGATTTTTCATGCGCATGGCC |
| | MatB-F189M-V192C-R197C-FOR | GCTTACATTGCGTGATATGTGGCGTTGCACCGCAGGCGATTGCCTGATTCATGCGTTACCGATTTTTCATACACATGGCC |
| | MatB-F189M-V192C-T207A-FOR | GCTTACATTGCGTGATATGTGGCGTTGCACCGCAGGCGATCGTCTGATTCATGCGTTACCGATTTTTCATGCGCATGGCC |
| | MatB-F189M-R197C-T207A-FOR | GCTTACATTGCGTGATATGTGGCGTGTAACCGCAGGCGATTGCCTGATTCATGCGTTACCGATTTTTCATGCGCATGGCC |
| | MatB-V192C-R197C-T207A-FOR | GCTTACATTGCGTGATTTCTGGCGTTGCACCGCAGGCGATTGCCTGATTCATGCGTTACCGATTTTTCATGCGCATGGCC |
| | MatB-F189M-V192C-R197C-T207A-FOR | GCTTACATTGCGTGATATGTGGCGTTGCACCGCAGGCGATTGCCTGATTCATGCGTTACCGATTTTTCATGCGCATGGCC |
| | MatB-T207S-FOR | GCTTACATTGCGTGATTTCTGGCGTGTAACCGCAGGCGATCGTCTGATTCATGCGTTACCGATTTTTCATAGCCATGGCC |
| | MatB-F189M-T207S-FOR | GCTTACATTGCGTGATATGTGGCGTGTAACCGCAGGCGATCGTCTGATTCATGCGTTACCGATTTTTCATAGCCATGGCC |
| | MatB-V192C-T207S-FOR | GCTTACATTGCGTGATTTCTGGCGTTGCACCGCAGGCGATCGTCTGATTCATGCGTTACCGATTTTTCATAGCCATGGCC |
| | MatB-R197C-T207S-FOR | GCTTACATTGCGTGATTTCTGGCGTGTAACCGCAGGCGATTGCCTGATTCATGCGTTACCGATTTTTCATAGCCATGGCC |
| | MatB-F189M-V192C-T207S-FOR | GCTTACATTGCGTGATATGTGGCGTTGCACCGCAGGCGATCGTCTGATTCATGCGTTACCGATTTTTCATAGCCATGGCC |
| | MatB-F189M-R197C-T207S-FOR | GCTTACATTGCGTGATATGTGGCGTGTAACCGCAGGCGATTGCCTGATTCATGCGTTACCGATTTTTCATAGCCATGGCC |
| | MatB-V192C-R197C-T207S-FOR | GCTTACATTGCGTGATTTCTGGCGTTGCACCGCAGGCGATTGCCTGATTCATGCGTTACCGATTTTTCATAGCCATGGCC |
| | MatB-F189M-V192C-R197C-T207S-FOR | GCTTACATTGCGTGATATGTGGCGTTGCACCGCAGGCGATTGCCTGATTCATGCGTTACCGATTTTTCATAGCCATGGCC |
| | MatB-P232L-REV | GCATGGTAGCCTGCGGCATCAAAGACAGGATCTCTTCCAGATCAAATTTGCTTAACAG |

| | | |
|---|---|---|
| | MatB-E233V-REV | GCATGGTAGCCTGCGGCATCAAAGACAGGATCTCCACCGGATCAAATTTGCTTAACAG |
| | MatB-P232L-E233V-REV | GCATGGTAGCCTGCGGCATCAAAGACAGGATCTCCACCAGATCAAATTTGCTTAACAG |
| 2 | MatB-H293S-M306-FOR | GAGATCCTGTCTTTGATGCCGCAGGCTACCATGCTGATGGGCG |
| | MatB-H293S-REV | CATACGGATTTGAGGTGTTCATGTTGGTCTCGGTCATTCCGTAGCGTTCCAGAATGGCGCTGCCGGTACGC |
| | MatB-M306I-REV | CATACGGATTTGAGGTGTTAATGTTGGTCTCGGTCATTCCGTAGCGTTCCAGAATGGCATGGCCGGTACGC |
| | MatB-M306V-REV | CATACGGATTTGAGGTGTTCACGTTGGTCTCGGTCATTCCGTAGCGTTCCAGAATGGCATGGCCGGTACGC |
| | MatB-H293S-M306I-REV | CATACGGATTTGAGGTGTTAATGTTGGTCTCGGTCATTCCGTAGCGTTCCAGAATGGCGCTGCCGGTACGC |
| | MatB-H293S-M306V-REV | CATACGGATTTGAGGTGTTCACGTTGGTCTCGGTCATTCCGTAGCGTTCCAGAATGGCGCTGCCGGTACGC |
| 3 | MatB-Combi-FOR | AACACCTCAAATCCGTATGAGGG |
| | MatB-Combi-REV | ATCACGCAATGTAAGCGCG |
